# The human cytomegalovirus chemokine binding protein UL22A is necessary for efficient reactivation from latency in CD34^+^ hematopoietic progenitor cells and humanized mice

**DOI:** 10.64898/2025.12.04.692306

**Authors:** Rebekah L Turner, Nicole L Diggins, Luke Slind, Jennifer Mitchell, Andrew H Pham, Christopher J Parkins, Wilma Perez, Samuel Medica, Michael Denton, Takeshi F Andoh, Gabriela M Webb, Daniel Andrade-Vera, Daniel N Streblow, Patrizia Caposio, Meaghan H Hancock

## Abstract

Herpesviruses and Poxviruses encode secreted chemokine binding proteins that prevent the interaction between chemokines and their cognate G protein coupled receptors to alter chemotactic gradients and intracellular signaling pathways. Human cytomegalovirus (HCMV) encodes the secreted protein UL22A (formerly UL21.5), which is described as a CCL5 (RANTES) binding protein and requires sulfation at two tyrosine residues (Y65 and Y69) for efficient RANTES interaction. In this report, we show that the UL22A protein, and the UL22A Y65 and Y69 residues are necessary for efficient HCMV reactivation from latency in CD34^+^ hematopoietic progenitor cells and that UL22A expression is essential for reactivation in a humanized mouse model of latency. However, RANTES neutralization is not sufficient to complement the *in vitro* reactivation defect of UL22A mutant viruses. These data suggest that UL22A plays an important role in latency, possibly through interactions with additional chemokines or other types of ligands via its tyrosine residues, in order to mediate efficient HCMV reactivation.

**IMPORTANCE:** HCMV is a ubiquitous herpesvirus that infects 60-90% of the population worldwide. In immunocompetent individuals, primary infection is asymptomatic and results in lifelong latent infection in CD34^+^ hematopoietic progenitor cells (HPCs). Viral reactivation remains a major complication for immunosuppressed individuals, but current therapeutics targeting HCMV replication show significant toxicity. Thus, a better understanding of the mechanisms controlling latency and reactivation is necessary to develop new therapeutics targeting these stages of the HCMV lifecycle. We show that virus lacking the HCMV chemokine binding protein UL22A is incapable of efficient reactivation in CD34^+^ HPCs and *in vivo*. UL22A tyrosine residues important for interaction with the chemokine RANTES are necessary for reactivation. However, neutralizing RANTES does not complement the reactivation defect of UL22A mutant viruses, demonstrating that UL22A has functions other than RANTES binding. Together, our results reveal a novel role for UL22A in HPCs and a new understanding of UL22A-chemokine interactions.

## INTRODUCTION

Chemokines are chemotactic cytokines expressed by cells in response to homeostatic or inflammatory stimuli that can direct the migration of leukocytes to specific sites and regulate their effector functions (1). Chemokines are structurally related proteins divided into four families (C, CC, CXC and CX3C) based on the spacing of cysteine residues and disulphide bridges that subtly determine chemokine structure. Chemokines interact with both surface glycosaminoglycans (GAGs), which is essential for the formation of immobilized chemotactic gradients that help guide cellular migration. Chemokine interaction with G protein-coupled receptors (GPCRs) is key to inducing cell signaling pathways involved in cytoskeletal rearrangements, adhesion, and extravasation. Chemokine signaling networks are incredibly plastic; most chemokine receptors bind multiple chemokines, and individual chemokines can bind to multiple receptors (2). This plasticity allows for fine tuning of the complex signaling patterns resulting in distinct functional outcomes depending on the chemokine milieu and the cell types involved. Given the important role of chemokines in the defense against pathogens, chemokines and chemokine receptor signaling pathways are important targets for virus interference (3, 4). Chemokine binding proteins (CKBP) encoded by herpesviruses and poxviruses demonstrate promiscuous binding to cellular chemokines; often binding to multiple chemokines within a single family, binding both chemokine and non-chemokine proteins or, in the case of murine gammaherpesvirus 68 (MHV68) M3 protein, binding to chemokines from all four families (5). Interestingly, CKBPs bear little resemblance to GPCRs, have no mammalian homologues, and show little structural homology to each other. However, the wide-ranging binding capacity of CKBPs allows viruses to efficiently interfere with chemokine function using little genetic material.

Human cytomegalovirus (HCMV) is a member of the betaherpesvirus family that causes lifelong persistence in the host through the establishment of latent infections in CD34^+^ hematopoietic progenitor cells (HPCs) and CD14^+^ monocytes (6). Reactivation from latency occurs via mechanisms that remain poorly defined, but immunosuppression, as during solid organ or hematopoietic stem cell transplantation, results in widespread virus replication and significant morbidity and mortality (7). Furthermore, congenital HCMV infection, occurring due to primary infection or virus reactivation during pregnancy, remains the leading cause of sensorineural defects in newborns (8). Understanding the mechanisms that control HCMV latency and reactivation are critical for developing new therapeutics to limit the effects of HCMV in these vulnerable populations.

The long co-evolution of HCMV with humans has resulted in the generation of sophisticated immune evasion mechanisms, including dysregulation of chemokine networks. HCMV disrupts chemokine function by encoding viral chemokines (e.g., UL128 and UL146 (9–11)), chemokine receptors (e.g., US28 (12–14)), regulating expression of cellular chemokines (15–18) and their receptors (19–22) and encoding the CKBP UL22A (23). UL22A is an abundant, secreted protein that is not essential for lytic infection of fibroblasts or endothelial cells. An initial study of UL22A tested its ability to interact with 3 of the approximately 45 known human chemokines and found it only bound to the CC chemokine CCL5 (regulated on activation, normal T cell expressed and secreted; RANTES) to prevent its interaction with the cell surface (23). Thus, the only known function of UL22A is as a RANTES binding protein, and the importance of this interaction to viral pathogenesis is still unclear.

Almost half of all cellular chemokine receptors contain sulfated tyrosines, which are important to enhance the interaction between the chemokine and its receptor (24). Sulfated tyrosines are often found within flexible anionic regions in chemokine receptors where they can make electrostatic interactions with chemokines at basic residues as well as through the tyrosine aromatic ring. Indeed, the HCMV GPCR US28 contains a tyrosine residue at position 16 which is essential for chemokine binding and is postulated to be sulfated (25). Previous work identified two tyrosine residues in the UL22A sequence, which when sulfated, enhanced the interaction between UL22A and RANTES, identifying UL22A as the first viral CKBP to harbor sulfated tyrosines (26).

In this report, we show that UL22A, while non-essential for lytic replication, plays an important role during latency in CD34^+^ HPCs and humanized mice and that two tyrosine residues in UL22A characterized as sulfated are essential for efficient reactivation from latency. However, complementing the only known function of UL22A, through neutralization of RANTES during HPC infection, does not restore the reactivation defect observed for HCMV UL22A mutants. Thus, the RANTES binding function of UL22A is not sufficient for promoting viral reactivation from latency, and suggests that the neutralization of other UL22A binding partners and/or yet-defined functions of UL22A requiring sulfated tyrosine residues are critical for HCMV reactivation.

## RESULTS

### UL22A is an early-late protein not essential for lytic replication

Early studies of UL22A investigated its functions in the context of the AD169 and VR1814 strains of HCMV (23). To investigate the role of UL22A in latent infection of CD34^+^ HPCs, we generated recombinant viruses based on the TB40/E strain of HCMV that contained an N-terminal HiBiT tag (encoded after the predicted UL22A signal sequence) and a C-terminal FLAG epitope (UL22AHF). In addition, we engineered a recombinant virus based on this construct that contains two contiguous stop codons immediately after the initiating methionine codon termed UL22AmutHF. We assessed the growth kinetics of TB40/E UL22AHF and UL22AmutHF in both single and multi-step growth curves and did not detect any defect in replication in the absence of UL22A (Fig 1A-D), as has been previously reported for HCMV strain AD169. UL22A was originally described as a spliced late gene (27) but was later reclassified as an early-late gene during lytic infection (28). However, the kinetics of protein expression has never been reported. We performed an analysis of UL22A protein expression upon infection of human fibroblasts in the presence or absence of the viral DNA polymerase inhibitor Foscarnet. Normal human dermal fibroblasts (NHDF) were mock infected or infected with UL22AHF or UL22AmutHF at a multiplicity of infection (MOI) of 3 and protein lysates were harvested from 8 to 72 hours post-infection. As shown in Fig 1E, immunoblotting using an antibody to the FLAG epitope demonstrated protein detection starting at 48 hours post-infection in cells infected with the UL22AHF virus, but not in cells infected with UL22AmutHF. Furthermore, protein was still detected, albeit at lower levels, in the presence of Foscarnet. We also took advantage of the highly sensitive detection capacity of the HiBiT epitope to quantitate UL22A secretion into the supernatants of the same infected cultures. As shown in Fig 1F, UL22A is detected in the supernatant as early as 24 hours post-infection, with only moderate effects of Foscarnet treatment. Thus, in the context of the TB40/E strain of HCMV, UL22A is a non-essential, early-late secreted protein that can be sensitively detected by the HiBiT tag.

**Figure 1.**
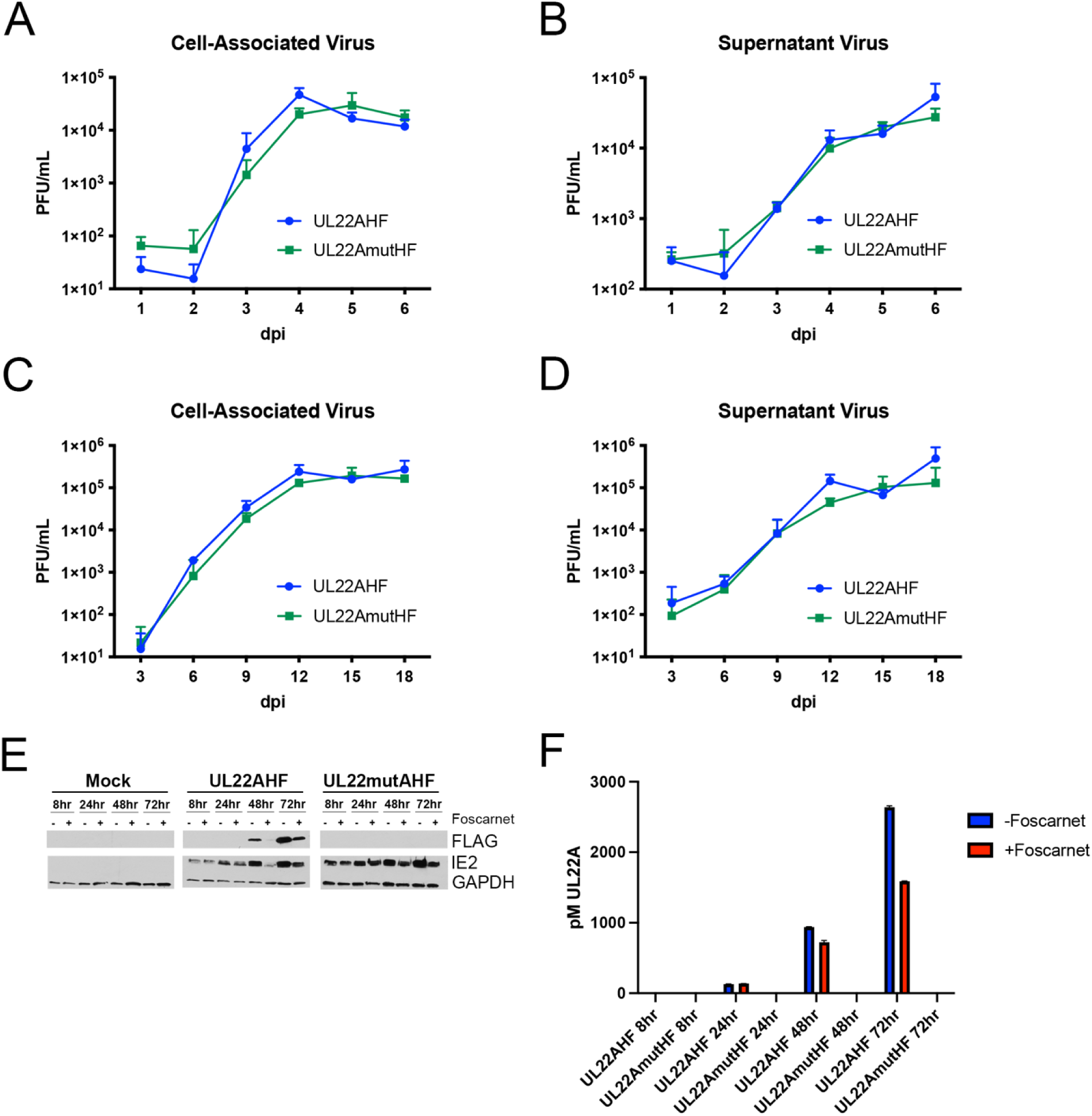
UL22A is an early-late secreted protein not necessary for viral replication. NHDFs were infected with TB40/E UL22AHF or TB40/E UL22AmutHF at an MOI of 3 for single step (A and B) or an MOI of 0.01 for multistep (C and D) growth curves. Cell associated and supernatant virus were harvested at the indicated timepoints, and PFU/mL values were quantified by titration on NHDF. n=3 (E and F) NHDFs were infected with TB40/E UL22AHF or TB40/E UL22AmutHF at an MOI of 3 and treated +/- foscarnet (200ug/mL). Cell lysates (E) and supernatants (F) were harvested at the indicated timepoints. (E) Cell lysates were immunoblotted for FLAG, IE2, and GAPDH. (F) Cell supernatants were quantified using the Nano-Glo Extracellular Detection System with HiBiT protein standard.

### UL22A is necessary for efficient reactivation from latency in CD34^+^ HPCs and humanized mice

UL22A transcripts are amongst the highest expressed in infected HPCs (29, 30), but a role for UL22A during infection of hematopoietic cells has not been reported. Thus, we next assessed the expression of UL22A during infection of CD34^+^ HPCs using the HiBiT detection assay. hESC-derived CD34^+^ HPCs were infected with TB40/E-GFP UL22AHF or UL22AmutHF at an MOI of 2 for 48 hours, and viable, CD34^+^, GFP^+^ cells were sorted and seeded into long-term bone marrow culture (LTBMC) over stromal cell support to allow for the establishment of latent infection, as previously described (31–33). At both 5 and 12 days in latency culture, supernatants were harvested and UL22A protein was assayed via the HiBiT epitope. As shown in Fig. 2A, UL22A protein is detectable at both 5 and 12 days in latency culture, with protein accumulating over time during infection with UL22AHF. As expected, the protein was not detected during infection with HCMV UL22AmutHF.

**Figure 2.**
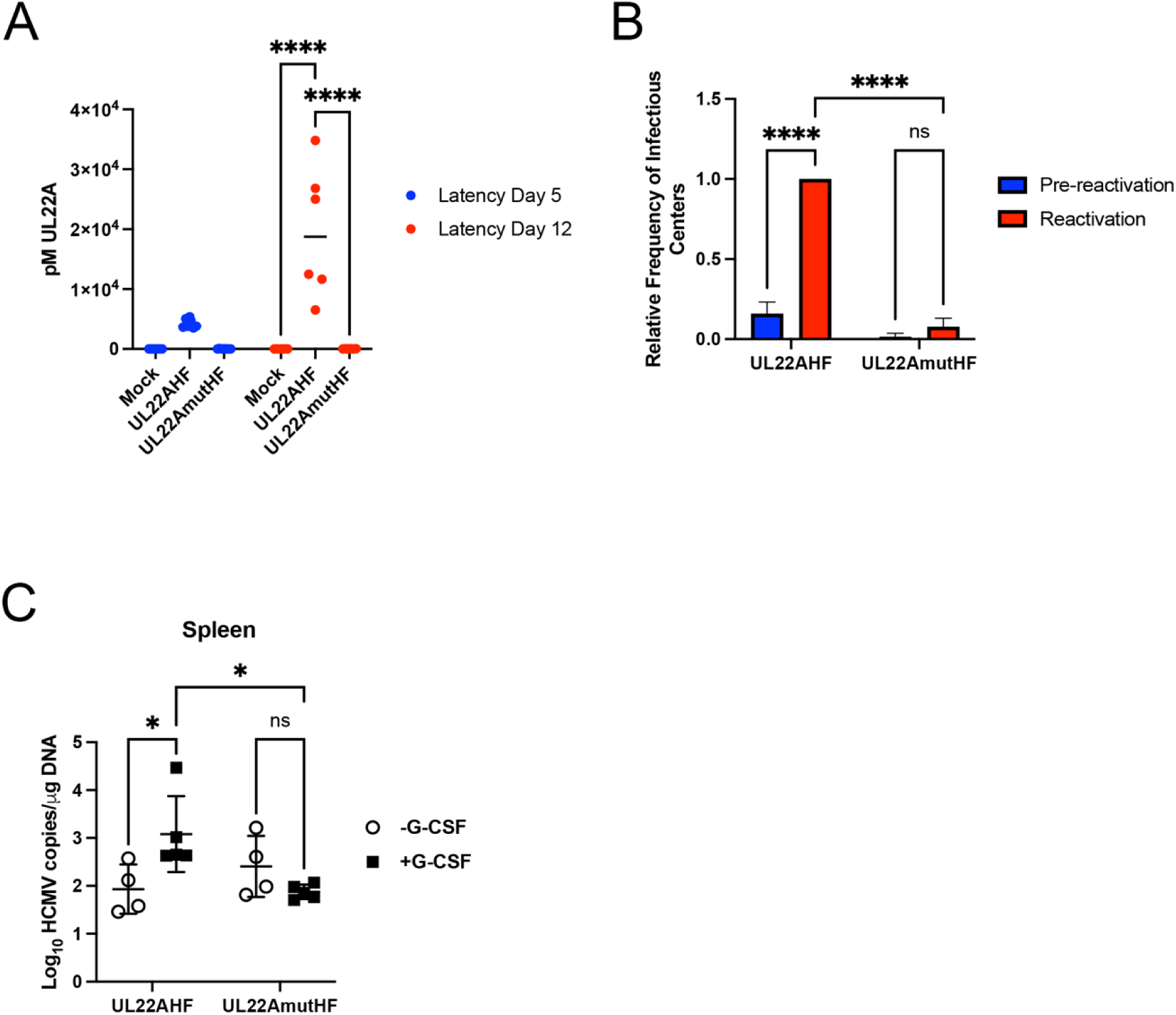
UL22A is secreted during latent infection and is necessary for reactivation from latency. (A and B) hESC-derived CD34^+^ HPCs were infected with TB40/E UL22AHF or UL22AmutHF at an MOI of 2 for 48 hours and then sorted using FACS for viable, CD34^+^, and GFP^+^ cells. For 12 days, cells were maintained in LTBMC in transwells above a murine stromal support to establish latent infection. (A) Supernatants were harvested at days 5 and 12 of latency culture. UL22A was detected using the Nano-Glo HiBit Extracellular Detection System (****p<0.0001 [two-way ANOVA with Tukey’s multiple comparison test]; n=6). (B) At 14 dpi, an equal number of cells were mechanically disrupted and seeded over a fibroblast monolayer to measure virus in latency culture (pre-reactivation) or cultured in cytokine-rich media to measure reactivation. At 21 days post-plating, GFP^+^ wells were counted, and the frequency of infectious centers were determined by ELDA software (****p<0.0001 [two-way ANOVA with Tukey’s multiple comparison test]; n=3). (C) Sub-lethally irradiated NOD-*scid* IL2Rγ_c_^null^ mice were engrafted with CD34^+^ HPCs (huNSG) and subsequently injected with human fibroblasts previously infected with TB40/E UL22AHF or UL22AmutHF (n=5). At 4 weeks post-infection, mice were treated with G-CSF and AMD3100 to promote viral reactivation. At 1-week post-treatment, mice were euthanized, and tissue was harvested. Total genomic DNA was isolated from splenic tissue, and HCMV genomes were quantified using qPCR with primers and probes specific for the UL141 gene (*p<0.05 [two-way ANOVA followed by Bonferroni’s multiple comparison test]).

The ability of the UL22AHF and UL22AmutHF recombinant viruses to reactivate from latency was evaluated in the hESC-derived CD34^+^ HPCs. After 12 days of LTBMC, HPCs were stimulated to differentiate and re-initiate virus replication by seeding onto monolayers of permissive fibroblasts in an extreme limiting dilution assay (ELDA; (34)) in cytokine-rich media. An equivalent number of cells were mechanically lysed and plated similarly to measure the production of free infectious virus during latency culture and serve as a pre-reactivation control. As shown in Fig. 2B, HPCs infected with UL22AmutHF reactivate virus at a lower frequency compared to cells infected with UL22AHF. Similar results were observed in fetal liver-derived CD34^+^ HPCs (Fig. S1). These findings indicate that the HCMV CKBP UL22A plays an important role in latency and/or reactivation in CD34^+^ HPCs *in vitro*.

The importance of UL22A in the context of an *in vivo* infection was investigated in the humanized mouse model of HCMV latency and reactivation (35, 36). In this model, CD34^+^ HPCs were engrafted into NSG mice. Following an eight-week period, the animals were infected by intraperitoneal injection with WT or UL22A mutant virus-infected fibroblasts. Four weeks later, mice were treated with G-CSF and AMD3100 to induce mobilization of myeloid cells from the bone marrow and promote viral reactivation (36). In mice infected with wild-type HCMV, the G-CSF/AM3100 mobilization triggered a marked increase in viral DNA copy number detected in the spleen (Fig. 2C), consistent with efficient reactivation. In contrast, mice infected with the UL22A mutant virus failed to exhibit a significant increase in viral DNA following mobilization. These *in vivo* findings recapitulate the *in vitro* results and suggest that UL22A is also essential for HCMV reactivation from latency in both cultured CD34^+^ HPCs and humanized mice.

### Generation and characterization of UL22A mutants that cannot interact with RANTES

Sulfation of tyrosine residues within the N-terminal extracellular domain of chemokine receptors increases their affinity for chemokines, enhancing their binding and functional activities (24). Previous work determined that UL22A harbors tyrosine residues that are subject to sulfation (26). UL22A proteins chemically synthesized to be singly or doubly sulfated were able to more efficiently compete RANTES from a CCR3 peptide compared to unsulfated protein, indicating that tyrosine sulfation plays an important role in UL22A’s interaction with RANTES. We validated these observations in a biological setting, whereby we generated plasmids expressing a nanoluciferase Large BiT (LgBiT) fusion-tagged UL22A where tyrosines at positions 65 and/or 69 were replaced with phenylalanine residues (Y65F, Y69F, Y65F+Y69F). We also generated MHV68 M3 tagged with LgBiT as a positive control, and the secreted HCMV protein UL7 tagged with LgBiT as a negative control. To measure interactions with RANTES, an additional plasmid was generated whereby RANTES was tagged with the complementing fragment of the nanoluciferase enzyme Small BiT (SmBiT) (Fig. 3A). We chose to use SmBiT as the complementing component of the nano-luciferase enzyme as it provides the optimal balance between interaction affinity and signal strength (37). IL8 and MCP-1 plasmids tagged with SmBiT were also cloned to characterize the specificity of interactions with UL22A, UL7 and M3. To investigate the interactions between the CKBPs and chemokine proteins, HEK293 cells were transfected with each LgBiT construct. 48 hours post-transfection supernatants were harvested and assayed for LgBiT protein expression using a bioluminescent Lumit assay and a HiBiT control protein. Equivalent amounts of UL22A, M3 or UL7 protein were then mixed with equivalent amounts of supernatant from HEK293 cells transfected with either RANTES-, IL8- or MCP-1-SmBiT. Binding interactions between the proteins in solution, resulting in an intact luciferase protein, was measured by bioluminescence assay. As shown in Fig. 3B, M3 interacts with RANTES, IL8 and MCP-1, as has previously been described (38, 39) while UL22A interacts with RANTES but not IL8 or MCP-1 (23, 26). In contrast, UL7 showed no detectable interaction with RANTES, IL8, or MCP-1 under the same conditions. However, UL22A constructs containing amino acid substitutions Y65F and/or Y69F were unable to interact with RANTES as efficiently as WT UL22A (Fig. 3C) nor did they interact with IL8 or MCP-1.

**Figure 3.**
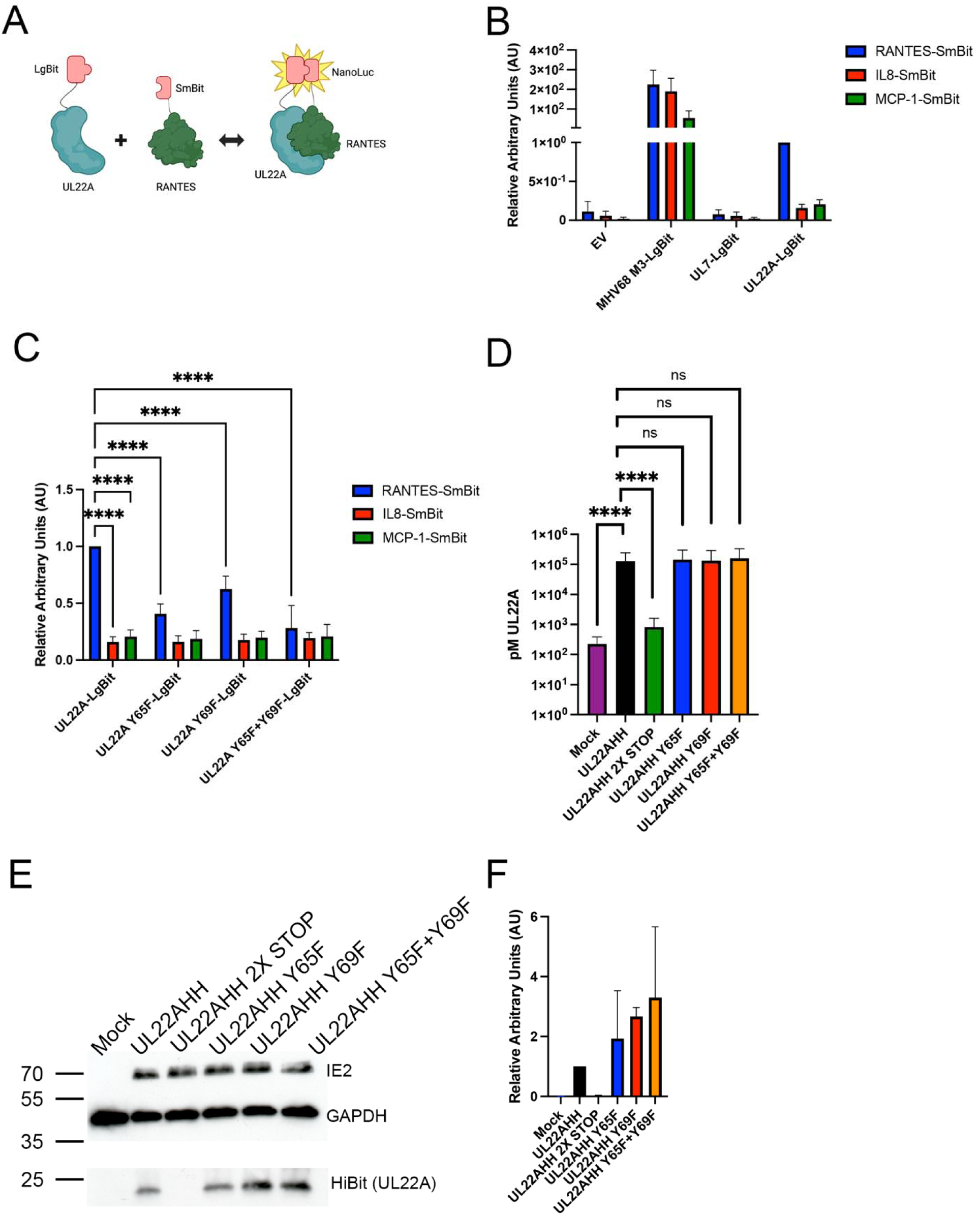
UL22A tyrosine residues Y65 and Y69 are important for binding RANTES. (A) Schematic of the nano-luciferase assay for measuring UL22A-RANTES interactions in solution. (B,C) HEK293 cells were transfected with 1.6μg pcDNA3.1-empty vector (EV), MHV68 M3-LgBit, UL7-LgBit, UL22A-LgBit, RANTES-SmBit, IL8-SmBit, or MCP-1-SmBit or (C) UL22A-LgBit, UL22A Y65F-LgBit, UL22A Y69F-LgBit, UL22A Y65+Y69F-LgBit, RANTES-SmBit, IL8-SmBit or MCP-1 SmBit. At 48 hours post-transfection, supernatants were harvested, LgBit proteins and SmBit proteins were normalized and equivalent amount EV or LgBit supernatants and equivalent amounts of SmBit supernatants were mixed. LgBit-SmBit interaction was quantified using the Nano-Glo Extracellular Detection Kit (Promega). (****p<0.0001 [two-way ANOVA with Tukey’s multiple comparison test]; n=3). Cell supernatants (D) and cell lysates (E) were harvested at 2dpi. (D) HiBit-tagged UL22A in supernatant was quantified across 3 independent experiments using the Nano-Glo Extracellular Detection System with HiBit protein standard (****p<0.0001 [unpaired t test]). (E) Cell lysates were immunoblotted for HiBit (UL22A), IE2, and GAPDH and 3 representative blots were quantified (F) using ImageJ software.

To evaluate the importance of UL22A RANTES binding in reactivation from latency, we derived TB40/E mutants containing the UL22A substitutions Y65F and Y69F using a recombinant virus expressing UL22A with both HiBiT and His epitopes at the C-terminus termed UL22AHH, UL22AHH Y65F, UL22AHH Y69F and UL22AHH Y65+Y69F respectively. A UL22A 2X STOP mutant containing two contiguous stop codons after the initiating methionine codon (UL22AHH 2XSTOP) was generated as a control virus. Growth kinetic analysis revealed that these mutant viruses replicated similarly to the WT UL22AHH virus (Fig. S2), and produced and secreted similar amounts of UL22A into the supernatant (Fig. 3D, E), indicating no deficiencies in expression or secretion of the mutated proteins.

### UL22A Y65 and Y69 residues of UL22A are necessary for efficient reactivation from latency

Using the HCMV recombinant viruses containing mutations in UL22A described above, we performed latency and reactivation assays in hESC-derived CD34^+^ HPCs as described in Figure 2. Firstly, we determined that WT and each of the tyrosine mutants secreted UL22A into the supernatant to similar levels after 12 days in latency culture (Fig. 4A), indicating no deficiencies in expression or secretion of the mutated UL22A proteins in CD34^+^ HPCs. Furthermore, we show that HCMV UL22AHH 2X STOP mutant has significantly reduced reactivation capacity compared to the parent UL22AHH virus (Fig. 4B), like what we observed with the UL22AmutHF (Fig. 2B). Additionally, substitution of a single tyrosine mutation (Y65F or Y69F) also reduced the reactivation efficiency of the UL22AHH virus, as did a combination of both mutations (Y65F+Y69F). Assessing viral genome copy numbers after 12 days in latency culture shows that these mutations did not affect maintenance of the viral genome or genome-containing cells (Fig. 4C). These data indicate that the tyrosine residues in UL22A that are subject to sulfation are important for HCMV latency and/or reactivation in CD34^+^ HPCs.

**Figure 4.**
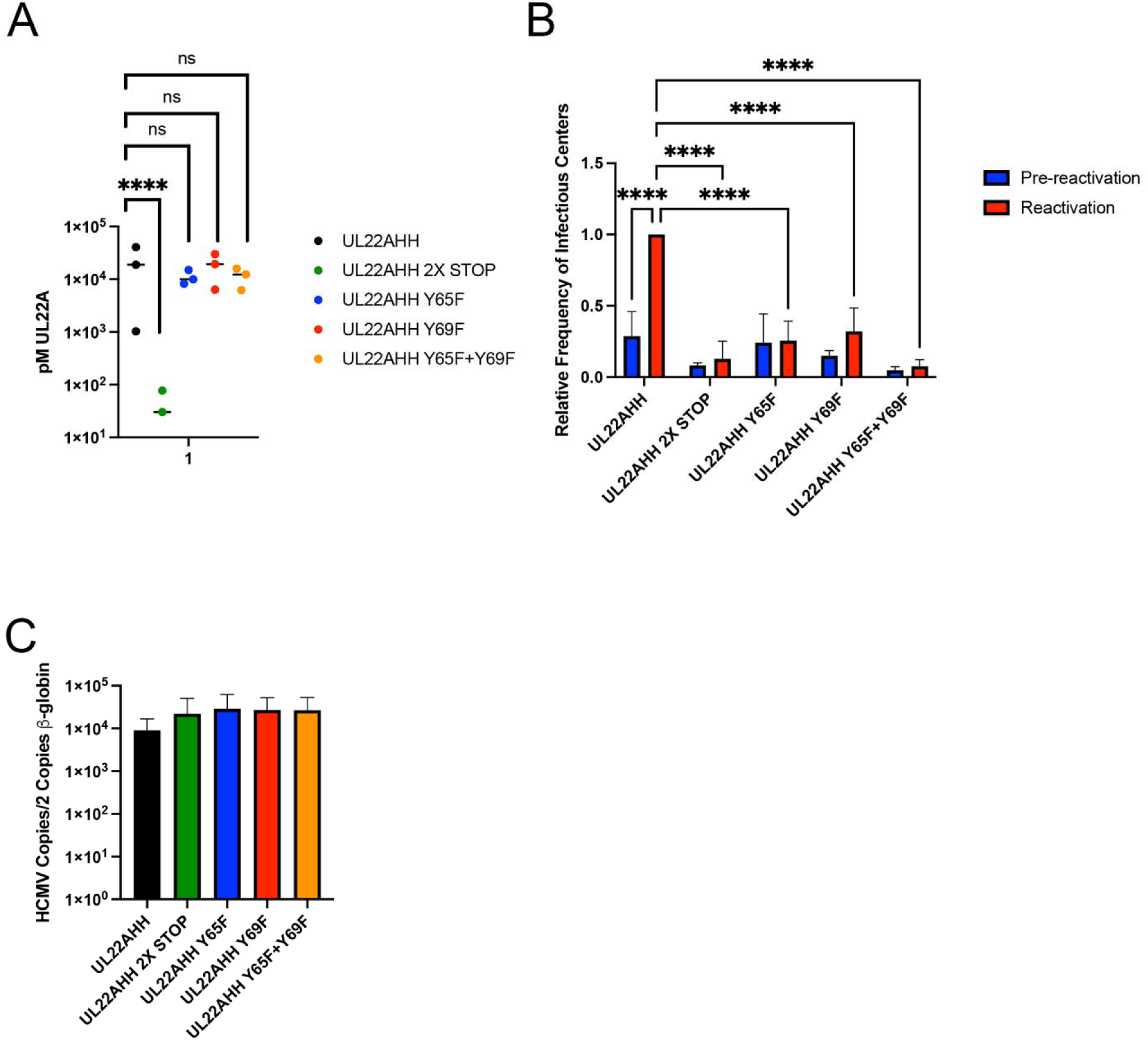
UL22A residues Y65 and Y69 are necessary for efficient reactivation from latency. hESC-derived CD34^+^ HPCs were infected with TB40/E UL22AHH, UL22AHH 2X STOP, UL22AHH Y65F, UL22AHH Y69F, UL22AHH Y65F+Y69F at an MOI of 2 for 48 hours and then sorted using FACS for viable, CD34^+^, GFP^+^ cells. (A) Sorted cells were put into latency culture as in Figure 2B. At 14dpi, supernatants were harvested from 3 independent experiments and HiBit-tagged UL22A in supernatant was quantified using the Nano-Glo Extracellular Detection System with HiBit protein standard (****p<0.0001 [with unpaired t test]; n=3). (B) Reactivation experiments were performed as in Figure 2B (****p<0.0001 [two-way ANOVA with Tukey’s multiple comparison test]). (C) Total genomic DNA was isolated from latently infected HPCs at 14dpi in three independent experiments. qPCR was used to quantify the ratio of viral genomes (copies of HCMV UL141) to cellular genomes (per two copies of human β-globin).

### Neutralization of RANTES does not complement a loss of UL22A during HCMV latency or reactivation in CD34^+^ HPCs

Above we demonstrate that expression of UL22A during HCMV infection in CD34^+^ HPCs, and more specifically both tyrosine residues, are necessary for efficient reactivation from latency. Thus, we hypothesized that UL22A binding to RANTES is a critical function that is required for viral reactivation from latency. To investigate whether the role of UL22A is to act as a decoy for RANTES by preventing interactions with chemokine receptors, we performed latency and reactivation assays in CD34^+^ HPCs that were infected with UL22AHH, UL22AHH 2X STOP and Y65F+Y69F in the presence or absence of RANTES neutralizing antibody. RANTES ELISA was performed to determine the amount of RANTES produced during latency culture, which ranged from ∼20-50pg/mL (Fig. S3A). From this data, the amount of anti-RANTES antibody to neutralize 50pg/mL of RANTES was determined to be 0.5-1μg/mL (Fig. S3B). Furthermore, 0.5-5μg/mL of anti-RANTES antibody inhibited RANTES-dependent calcium flux in CME cells (Fig. S3C). Thus, we performed latency and reactivation experiments as described above, including either IgG control antibody or RANTES neutralizing antibody at a concentration of 1μg/mL when cells were plated in latency, and the antibodies were replenished again after 5 days in latency culture. As is shown in Fig. 5A, addition of the IgG control antibody or the RANTES neutralizing antibody did not significantly affect the ability of the UL22AHH virus to reactivate from latency. Significantly, the RANTES neutralizing antibody was unable to complement the reactivation defect of either of the UL22AHH 2X STOP or UL22AHH Y65F+Y69F mutants, indicating that neutralization of RANTES is not the essential function of UL22A during latency in CD34^+^ HPCs. However, neutralization of RANTES during the reactivation process might be required to promote this process in an efficient manner. Therefore, reactivation assays were performed in cells infected with UL22AHH, UL22AHH 2X STOP and UL22AHH Y65F+Y69F mutants and the RANTES neutralizing and control antibodies were added only during the reactivation phase. The antibodies were replenished after 1 week following initiation of reactivation. As is shown in Fig. 5B, addition of IgG control or RANTES neutralizing antibody did not affect the reactivation capacity of the UL22AHH virus and similarly did not complement the reactivation defect of the UL22A 2X STOP or UL22AHH Y65F+Y69F mutants. Based on these findings, and to ensure our experimental conditions were fully neutralizing RANTES, reactivation assays using an excess of RANTES neutralizing antibody (5ug/mL) were also performed. This increased antibody concentration did not complement reactivation of the UL22AHH Y65+Y69F mutant (Fig. S4). Thus, these data suggests that while the tyrosine residues that are sulfated in UL22A are important for reactivation, RANTES binding may not be the only function of UL22A responsible for efficient HCMV reactivation from latency in CD34^+^ HPCs.

**Figure 5.**
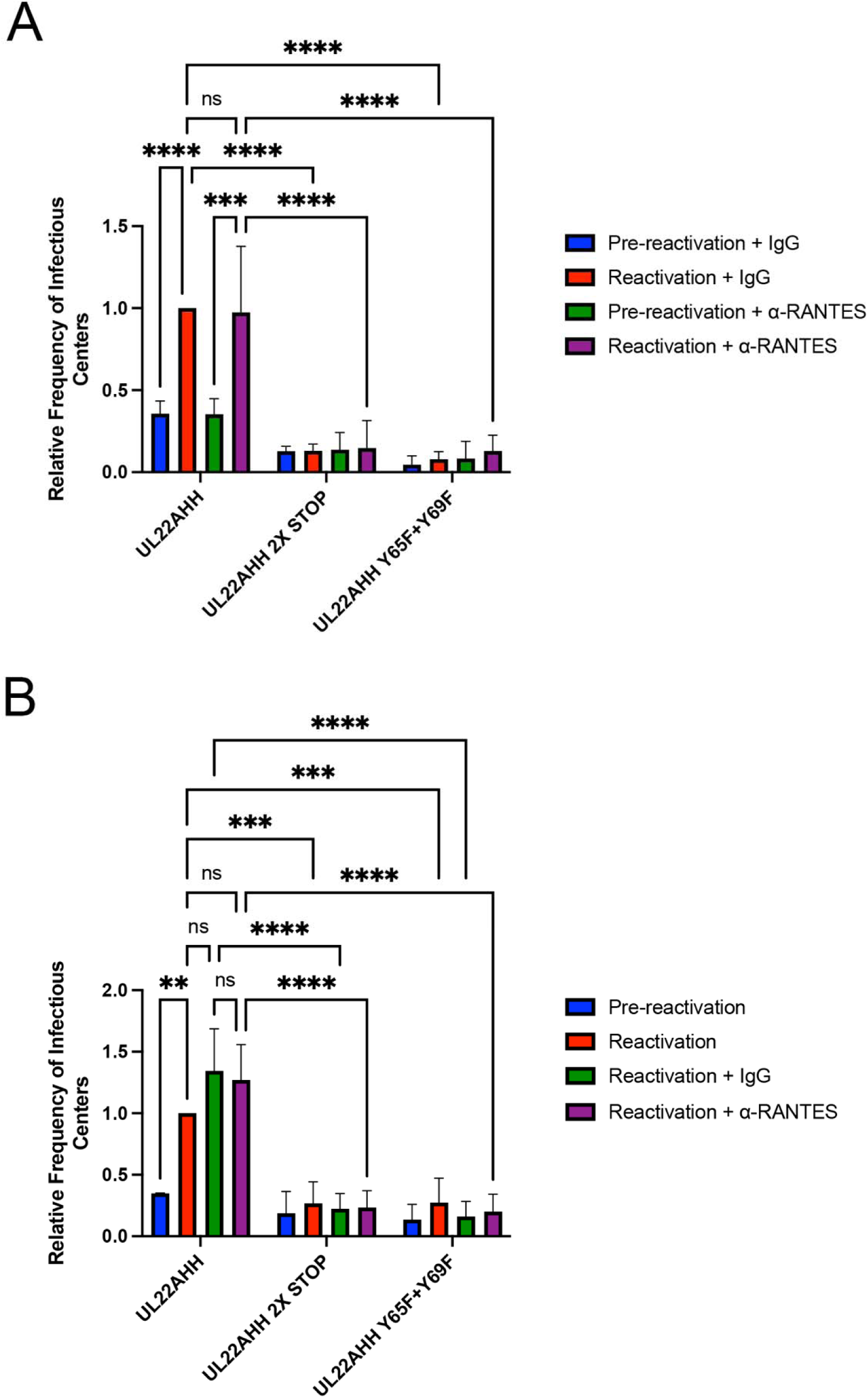
Neutralization of RANTES during latency or reactivation does not complement the reactivation defect of UL22A mutant viruses. hESC-derived CD34^+^ HPCs were infected with TB40/E UL22AHH, UL22AHH 2X STOP, UL22AHH Y65F, UL22AHH Y69F, UL22AHH Y65F+Y69F at an MOI of 2 for 48 hours and then sorted using FACS for viable, CD34^+,^ GFP^+^ cells. (A) Cells were treated with 1μg/mL of α-RANTES antibody or a control IgG and replenished again after 5 days while maintained in LTBMC in transwells above a murine stromal support to establish latent infection. (B) For 12 days, cells were maintained in LTBMC medium in transwells above a murine stromal support to establish latent infection. At 14 dpi, an equal number of cells were mechanically disrupted and seeded over a fibroblast monolayer to measure virus in latency culture (pre-reactivation) or cultured in cytokine-rich media containing 1μg/mL of α-RANTES antibody or a control IgG to measure reactivation. Control IgG or RANTES neutralizing antibody were replenished after 1 week of reactivation. At 21 days post-plating, GFP^+^ wells were counted, and the frequency of infectious centers were determined by ELDA software (**p<0.005, ***p<0.0005, ****p<0.0001 [two-way ANOVA with Tukey’s multiple comparison test]; n=3).

## DISCUSSION

The work presented here describes an essential role for the viral CKBP UL22A during HCMV reactivation in CD34^+^ HPCs. HCMV lacking UL22A was unable to efficiently reactivate, but genome copies present during latency were similar to UL22A-intact virus, indicating that the impact of UL22A is either on promoting reactivation itself or establishing a cellular environment during latency that is conducive to reactivation. Further investigation is required to identify the precise step in the reactivation process that UL22A impacts. UL22A has two tyrosine moieties that are required for RANTES binding (26), and herein we validated this property with a Lumit-based assay. Replacement of either of the tyrosine residues significantly reduced binding of RANTES to UL22A, as did dual substitution (Fig. 3). We also demonstrated that UL22A failed to bind the closely related chemokine MCP-1 (CCL2) or the more distantly related chemokine IL8 (CXCL8), further confirming the specificity of RANTES binding. Recombinant HCMV containing a UL22A 2X STOP mutation or substitutions of one or more of the UL22A tyrosine moieties did not display any replication defects in human primary fibroblasts. However, these viruses were crippled in their ability to reactivate. Thus, we surmised that UL22A sequestration of RANTES was required for viral reactivation. Nonetheless, neutralization of RANTES during latency or reactivation did not impact wild type virus reactivation nor did it restore the reactivation phenotype of viruses lacking functional critical residues for RANTES interaction. These findings suggest that functions mediated by UL22A other than, or in addition to, RANTES binding are required for efficient HCMV reactivation from latency.

While the UL22A mutant viruses (2X STOP and tyrosine mutants) are unable to efficiently reactivate from latency, viral genome copy numbers are similar between WT virus and the mutants (Fig. 4C), indicating that viral genomes, or genome-containing cells, are not lost during latency culture in the absence of UL22A function. However, it remains possible that UL22A may play a role in altering chemokine signaling during latency to ensure the viral genome is poised to respond to reactivation signals. Thus, we tested RANTES neutralization during both latency (Fig. 5A) and at the time of reactivation (Fig. 5B) and found that in neither case did addition of RANTES neutralizing antibody, at concentrations known to neutralize the upper limit of RANTES production during infection (Fig. S2) or in excess (Fig. S3), complement the reactivation defect of the UL22A mutants (Fig. 5). While we interpret this to mean that UL22A RANTES binding is not sufficient to stimulate virus reactivation, it remains possible that RANTES binding is one of several functions of UL22A, mediated by tyrosines 65 and 69, that is necessary to stimulate reactivation from latency.

The requirement for tyrosine residues within an anionic region of UL22A for efficient reactivation suggests that binding to additional chemokines, via sulfated tyrosine residues, may be an essential function of UL22A. It is important to note that we were unable to validate that the UL22A Y65 and Y69 residues were sulfated during ectopic expression or in the context of infection due to limitations in the commercially available reagents available to assess protein sulfation. It remains formally possible that other post-translational modifications, such as tyrosine phosphorylation, may play a role in UL22A function. Herpesfold predicts no specific structural conformation for UL22A, thus whether the CKBP interacts with RANTES and other chemokines via additional motifs is still unknown. Using the split-nanoluciferase approach to investigate chemokine binding uncovered that UL22A binding to RANTES is several orders of magnitude less efficient than the interaction of M3 and RANTES (Fig. 3B). Whether this indicates that UL22A is generally a weaker chemokine binder than M3, or that RANTES is not its main interactor, remains to be determined. Aside from binding additional chemokines, other roles for UL22A during latent infection can be envisioned. Some poxvirus CKBP also bind non-chemokine proteins, like the myxomavirus M-T7 protein, which binds IFNγ in addition to a broad range of C, CC and CXC chemokines (40, 41). Alternatively, as a secreted protein UL22A may itself interact with cell surface receptors to mediate intracellular signaling events, facilitated by the tyrosine residues. Moreover, while we and others have clearly identified UL22A as a secreted protein (23), we cannot rule out an intracellular function for the protein that is important for reactivation from latency. However, only secreted and transmembrane proteins are reported to be sulfated (42), thus, if the Y65 and Y69 residues are sulfated in the context of infection, this suggests that the secreted form of UL22A is required for reactivation. More experimentation is required to identify any additional UL22A interactors in order to better understand its function in CD34^+^ HPCs.

For hematopoietic stem and progenitor cells, chemokines regulate the hematopoietic niche by inducing both retention of HPCs and their quiescence. Stromal cell-derived factor-1 (SDF-1; CXCL12) was the first chemoattractant described for CD34^+^ HPCs which interacts with CXCR4 on the HPC surface to promote cell retention in the bone marrow as well as proliferation and survival (43–45). Human CD34^+^ HPCs express CXCR1, CXCR2 and CCR5 on the cell surface in addition to CXCR4 (46), indicating that these cells are responsive to numerous chemokines, including RANTES. Intriguingly, many CC chemokines are strong inducers of HPC mobilization and have inhibitory effects on myelopoiesis (47, 48). HCMV modulates differentiation along the myeloid lineage to stimulate virus reactivation in terminally differentiated macrophages (49–51). Thus, one function of UL22A in the context of HPC infection may be to bind chemokines (including RANTES) to modulate the differentiation of HPCs.

As potent chemokine binders, the CKBP of herpes- and poxviruses have been investigated for their roles in regulating chemotaxis *in vitro* and *in vivo*. Many CKBP block leukocyte migration and the calcium flux initiated by chemokine binding to cell surface receptors, but their effect on pathogenesis in small animal models has generally been modest. In the context of poxviruses, larger lesion sizes were observed in animals infected with viral mutants lacking CKBPs due to a greater influx of leukocytes (52–60). During MHV68 infection, knockout of the M3 protein resulted in fewer latently infected splenic B cells, but the phenotype could be reversed with depletion of CD8^+^ T cells, suggesting that M3 may function to block recruitment of T cells and allow the establishment of a latent reservoir (61, 62). The importance of UL22A in *in vitro* latency and reactivation assays suggests that its ability to block chemokine receptor binding and subsequent effector functions specifically in HPCs is an important facet of its function but does not preclude a role in disrupting chemotactic gradients *in vivo*. UL22A homologues are only encoded by primate CMVs, and thus the role of the CKBP in modulating leukocyte recruitment in response to chemokines has not been directly investigated *in vivo*. However, here we show that UL22A is necessary for virus reactivation in our humanized mouse model. Whether UL22A, like CKBPs from other viruses, binds chemokines from different species is unknown and thus whether in this system UL22A functionally interacts with chemokines produced by, and mediates interactions with, mouse and/or human cells remains to be investigated.

UL22A was originally characterized as the most abundantly produced transcript during lytic infection (27) and early efforts to characterize the latent transcriptome in CD14^+^ monocytes and CD34^+^ HPCs also noted UL22A as amongst the highest expressed transcripts (29, 30). Moreover, in acutely infected rhesus macaques, the rhesus CMV homologue of UL22A, Rh38.1, is amongst the most abundantly detected transcript in tissues including the lung and lymph nodes (63). Here we now show that UL22A protein is also abundantly produced during lytic and latent infection and plays a critical role in regulating reactivation from latency in CD34^+^ HPCs and humanized mice. The function(s) of UL22A necessary to mediate reactivation remain to be determined, as the full binding breadth of this chemokine binding protein awaits characterization.

## MATERIALS AND METHODS

### Cells and media

Feeder-free hESCs were obtained from WiCell (WA01-H1, hPCSC Reg identifier (ID) WAe001-A, NIH approval no. NIHhESC-10-0043). Cells were thawed and plated on Matrigel-coated six-well plates in complete mTeSR1 (Stem Cell Technologies). CD34^+^ HPCs were differentiated using a commercial feeder-free hematopoietic differentiation kit (STEMdiff Hematopoietic Kit, Stem Cell Technologies). HEK293 and adult normal human dermal fibroblasts (NHDF) were obtained from ATCC and cultured in Dulbecco’s modified Eagle’s medium (DMEM) supplemented with 10% heat-inactivated fetal bovine serum (FBS; Hyclone), 100 units/ml penicillin, 100 μg/ml streptomycin, and 100 μg/ml glutamine (Thermofisher). M2-10B4 and S1/S1 stromal cells were obtained from Stem Cell Technologies and maintained in DMEM with 10% FBS and penicillin, streptomycin, and glutamine as previously described. All cells were maintained at 37°C and 5% CO_2_.

### Viruses

HCMV viruses used in this study were derived from BAC-generated WT TB40/E expressing GFP from the SV40 promoter (64). HCMV TB40/E mutant viruses containing HiBiT and FLAG or His tags as well as point mutations to generate UL22A 2XSTOP and UL22A Y65F, Y69F or Y65/69F were generated by galactokinase (galK)-mediated recombination. For the TB40/E UL22AHF viruses, the galK gene replaced the UL22A ORF using the following primers: pUL22A KO galk F: AGATGTCGTCACCCAAGGTATTTAACGGCACACAGCCAGACGCGTTCGTCAGCAGCGACGCCGACAAGACCTCAGCCCTGTTGACAATTAATCATC and pUL22A KO galk R: TTTTAGAGCAAAACCTTACAGCTTTTTAATAAAAAACAAGGTAGTCAACATAATCGTTAACCCTTGGGGTCTGCTGCTCAGCAAAAGTTCGATTTA. In the second recombination step, galK was removed and replaced with a UL22A ORF containing a HiBiT tag after the UL22A signal sequence and a 3’ FLAG tag. This construct was generated as a plasmid and amplified using the following primers: pUL22A-tag insert F: AGATGTCGTCACCCAAGGTATTTAACGGCACACAGCCAGACGCGTTCGTCAGCAGCGACG CCGACAAGACCTCAGCATGGCTCGGAGGCTATGGATCTTGpUL22A-tag insert R: TTAGAGCAAAACCTTACAGCTTTTTAATAAAAAACAAGGTAGTCAACATAATCGTTA ACCCTTGGGGTCTGCTGTTACTTGTCGTCATCGTCTTTGTAGT. The UL22AHFmut virus was generated using the same galK intermediate and incorporating 2 stop codons after the initiating methionine in the UL22AHF construct in the F primer: pUL22Amut-tag insert F: AGATGTCGTCACCCAAGGTATTTAACGGCACACAGCCAGACGCGTTCGTCAGCAGCGACGCCGACAAGACCTCAGCTAGGCTTAGAGGCTATGAATCTT and the same pUL22A-tag insert R as above. To generate the UL22AHH viral recombinant, we replaced a portion of the UL22A gene with galK using the following primers: UL22A 3 prime HiBiT/ galk F: CAGCGAACATCAGCAACCACAAAAGACTGATGAACACAAAGAAAATCAAGCCAAAGAAAATGAAAAGAAGATTCAG CCTGTTGACAATTAATCATCGGCA, UL22A 3 prime HiBiT/ galk R: TAGAGCAAAACCTTACAGCTTTTTAATAAAAAACAAGGTAGTCAACATAATCGTTAACCCTTGGGGTCTGCTGTTA CTCAGCAAAAGTTCGATTTA. The second step recombination product was generated by amplifying a portion of the UL22A gene from a plasmid encoding UL22A with a 3’ HiBiT and HIS tag (see *Reagents*) using the following primers: UL22A plasmid HiBiT amp F: GAACGCGGTTATCGTGTTTTTGCAGCGTGACGGTGGAACAACCCAGTACCAGCGCTGATGGTAGTAATACCACCCCCAGCAAGAACGTAACTCTCAGTCA and UL22A plasmid HiBiT amp R: TAGAGCAAAACCTTACAGCTTTTTAATAAAAAACAAGGTAGTCAACATAATCGTTAACCCTTGGGGTCTGCTGTTAGCTAATCTTCTTGAACAGCCGCCA. To generate the UL22AHH 2X STOP mutant, galK was inserted immediately after the UL22A stop codon using the following primers: UL22A mut galk F: TGTCGTCACCCAAGGTATTTAACGGCACACAGCCAGACGCGTTCGTCAGCAGCGACGCCGACAAGACCTCAGCATG CCTGTTGACAATTAATCATCGGCA, UL22A mut galk R: TCGATTTCTGAGAAGGTGCCGCCAAAGCCACCGTCAAGGTCACGGCTAGTAAGCTCAAGATCCATAGCCTCCGAGC CTCAGCAAAAGTTCGATTTA. 2 contiguous STOP codons were inserted following the initiating methionine using the following oligos: UL22A 2X stop oligo F: CAGCCAGACGCGTTCGTCAGCAGCGACGCCGACAAGACCTCAGCATGTAATAGAGGCTATGGATCTTGAGCTTACTAGCCGTGACCTTGACGGTGGCTTT, UL22A 2X stop oligo R: AAAGCCACCGTCAAGGTCACGGCTAGTAAGCTCAAGATCCATAGCCTCTATTACATGCTGAGGTCTTGTCGGCGTCGCTGCTGACGAACGCGTCTGGCTG. In order to generate UL22A mutants where tyrosine residues Y65 and Y69 were replaced with phenylalanine (Y65F, Y69F and Y65F + Y69F), the region encompassing these residues was replaced with the galK gene using the following primers: UL22A HiBiT His Y65/69F/ galk F: TGATGGTAGTAATACCACCCCCAGCAAGAACGTAACTCTCAGTCAGGGGGGGTCCACCACCGACGGAGACGAAGAT CCTGTTGACAATTAATCATCGGCA, UL22A HiBiT His Y65/69F/ galk R: CTTGATTTTCTTTGTGTTCATCAGTCTTTTGTGGTTGCTGATGTTCGCTGCCATCTCCGTCTGTAATCAAAACGTC CTCAGCAAAAGTTCGATTTA. For each tyrosine mutant, the following oligos were used to replace the galK insert: **Y65F:** UL22A HiBiT His Y65F oligo FACTCTCAGTCAGGGGGGGTCCACCACCGACGGAGACGAAGATTTTTCCGGGGAGTATGA CGTTTTGATTACAGACGGAGATGGCAGCGAACATCAGCAAC, UL22A HiBiT His Y65F oligo R: GTTGCTGATGTTCGCTGCCATCTCCGTCTGTAATCAAAACGTCATACTCCCCGGAAAAATCTTCGTCTCCGTCGGTGGTGGACCCCCCCTGACTGAGAGT. **Y69F:** UL22A HiBiT His Y69F oligo F: ACTCTCAGTCAGGGGGGGTCCACCACCGACGGAGACGAAGATTACTCCGGGGAGTTCGACGTTTTGATTACAGACGGAGATGGCAGCGAACATCAGCAAC, UL22A HiBiT His Y69 oligo R: GTTGCTGATGTTCGCTGCCATCTCCGTCTGTAATCAAAACGTCGAACTCCCCGGAGTAATCTTCGTCTCCGTCGGTGGTGGACCCCCCCTGACTGAGAGT. **Y65F + Y69F:** UL22A HiBiT His Y65F and Y69F oligo F: ACTCTCAGTCAGGGGGGGTCCACCACCGACGGAGACGAAGATTTTTCCGGGGAGTTCGACGTTTTGATTACAGACGGAGATGGCAGCGAACATCAGCAAC, UL22A HiBiT His Y65F and Y69F oligo R: GTTGCTGATGTTCGCTGCCATCTCCGTCTGTAATCAAAACGTCGAACTCCCCGGAAAAATCTTCGTCTCCGTCGGTGGTGGACCCCCCCTGACTGAGAGT.

All virus stocks were propagated and titered on NHDFs using standard techniques. To assess growth kinetics, NHDFs were infected at a MOI of 3 for single-step growth curves or a MOI of 0.01 for multi-step growth curves for 2 hr. Cell-associated and supernatant virus was harvested at multiple time points post-infection. Titers were determined by plaque assay on NHDFs.

### Reagents

The following commercial antibodies were used: GAPDH (ab8245, Abcam) and HCMV IE2 (MAB810, Sigma Aldrich). Neutralization of RANTES was accomplished with Human/Primate CCL5/RANTES antibody (MAB678, R&D Systems) and Mouse IgG_1_ isotype control antibody (MAB002, R&D Systems). HCMV UL22A was amplified by PCR with an upstream primer (5’-<u>GAATTC</u> GCCACCATGGCTCGGAGGCTATGGATC - 3’), and a downstream reverse primer (5’-<u>AAGCTT</u>TTATTAGCTAATCTTCTTGAACAGCCGCCAGCCGCTCACAGAGTGATGGTGATGGTGATGCTGAATCTTCTTTTC - 3’), containing EcoRI and HindIII restriction sites respectively, as well as HiBiT and His epitopes, and cloned into pcDNA3.1-with traditional cloning methods. HiBit-tagged proteins were visualized using the Nano-Glo HiBit Blotting System (N2410, Promega).

### Luciferase assays

For HiBit-tagged UL22A detection during viral infection, NHDFs were seeded in 12-well plates at a density of 1×10^6^ cells per plate and infected with indicated viruses at an MOI of 3 for 48 hours. Supernatants were harvested and amount of HiBit-tagged UL22A was determined using HiBit Control Protein (N3010, Promega) and a standard curve with the Nano-Glo HiBit Extracellular Detection Kit (Promega) according to the manufacturer’s protocol. To assess HiBit tagged UL22A during latent infection, supernatants were harvested at indicated timepoints and HiBit-tagged UL22A was determined as above.

For split-luciferase assays, 293 cells were seeded in 12-well plates at a density of 1.5×10^6^ per plate. The following day, 1.6ug of indicted LgBit and SmBit-tagged pcDNA3.1-constructs were transfected into separate wells using Lipofectamine 2000 (Invitrogen) following the manufacturer’s recommendation. At 48 hours post transfection, supernatants were harvested. In order to quantitate the amount of LgBit protein in the supernatants, 500pM HiBit Control Protein was mixed with the LgBit-tagged construct supernatants to generate a functional luciferase enzyme. As a control to measure background luminescence, 500pM HiBit Control Protein was added to 25uL of EV-transfected supernatants. To quantitate the amount of SmBit protein that was present in culture supernatants, purified LgBit protein was added to the supernatants of cells transfected with the SmBit-tagged constructs. Under these conditions, luciferase was measured using the Nano-Glo HiBit Extracellular Detection Kit according to the manufacturer’s protocol and total protein amounts were normalized. Following normalization, supernatants containing equivalent amounts LgBit protein or EV control were mixed with supernatant containing equivalent amounts of SmBit protein. LgBit-SmBit interaction was then measured using the Nano-Glo HiBit Extracellular Detection kit according to the manufacturer’s protocol. Luminescence was measured using a Promega GloMax Navigator Luminometer. Results were transferred to a Microsoft Excel spreadsheet and results were analyzed using GraphPad Prism 10 software.

### Ratiometric calcium flux measured by flow cytometry

CEM.NKR.CCR5 cells were washed with Hanks Balanced Salt Solution (HBSS; Invitrogen, Grand Island, NY, USA), resuspended at 1×10^7^ cells/ml in 37°C HBSS with 1uM Fura Red, AM (Invitrogen, Grand Island, NY, USA) an amine reactive dye for viability, and incubated in a 37°C water bath for 30 minutes. Cells were washed and resuspended in HEPES Buffered Saline Solution (HBSS with 1mM CaCl_2_, 0.5mM MgCl_2_, 0.1% BSA, 10mM HEPES) at 1×10^7^ cells/ml.

Recombinant RANTES (R&D Systems, Inc, MN, USA) and anti-RANTES antibodies were incubated at indicated concentrations for 30 minutes prior to use and were prepared at 10X the final concentration in HEPES Buffered Saline Solution. Background calcium flux was recorded for 20 seconds, sample was removed while recording, and 180μl of cell suspension was transferred to 20uL of RANTES (+/- anti-RANTES) (10x) and placed once again on the cytometer for recording. Events were recorded for a total of 120 seconds.

Ratiometric “Fura Red Ratio” was calculated as the increasing signal from the Violet laser (406 nm) over the decreasing signal from the Green laser (532 nm) using the Kinetics tool in FlowJo Software version 10.10.0 (TreeStar, Ashland, OR, USA). All flow cytometric data was collected using an LSR II (Becton Dickinson, Franklin Lakes, NJ, USA).

### Western blot analysis

Cells were harvested in protein lysis buffer (50mM Tris-HCl pH 8.0, 150mM NaCl, 1% NP40, and protease inhibitors), loading buffer (4X Laemmli Sample Buffer with 2-mercaptoethanol) was added, and lysates were incubated at 95°C for 5 min. Extracts were loaded onto 4-15% acrylamide gels (Biorad), transferred to Immobilon-P membranes (Millipore), and visualized with the specified antibodies. The relative intensity of bands detected by Western blotting was quantified using ImageJ software.

### CD34^+^ HPC latency and reactivation assays

CD34^+^ HPCs were infected with the indicated viruses at an MOI of 2 for 48hr, or were left uninfected, in stem cell media (Iscove’s modified Dulbecco’s medium [IMDM] [Invitrogen] containing 10% BIT serum replacement [Stem Cell Technologies], penicillin/streptomycin, stem cell factor [SCF], FLT3 ligand [FLT3L], interleukin-3 [IL-3], interleukin-6 [IL-6] [PeproTech], 50uM 2-mercaptoethanol, and 20ng/ml low-density lipoproteins). Pure populations of viable, infected (GFP^+^) CD34^+^ HPCs were isolated by fluorescence-activated cell sorting (FACS) (BD FACSAria equipped with 488-, 633-, and 405-nm lasers and running FACSDiva software) and used in latency assays as previously described (31–33). Briefly, cells were cultured in transwells above irradiated stromal cells (M2-10B4 and S1/S1) for 12 days to establish latency. Virus was reactivated by coculture with NHDF in RPMI medium containing 20% FBS, 1% P/S/G, and 15ng/ml each of G-CSF and GM-CSF in an extreme limiting dilution assay (ELDA). GFP^+^ wells were scored 3 weeks post-plating and the frequency of infectious centers was using ELDA software (34).

### Engraftment and infection of humanized mice

All animal studies were carried out in strict accordance with the recommendations of the American Association for Accreditation of Laboratory Animal Care. The protocol was approved by the Institutional Animal Care and Use Committee (protocol 0922) at Oregon Health and Science University. NOD-*scid*IL2Rγ_c_^null^ mice were maintained in a pathogen-free facility at Oregon Health and Science University in accordance with procedures approved by the Institutional Animal Care and Use Committee. Both sexes of animals were used. Humanized mice were generated as previously described (35, 36). The animals (12-14 weeks post-engraftment) were treated with 1 ml of 4% Thioglycollate (Brewer’s Media, BD) by intraperitoneal (IP) injection to recruit monocyte/macrophages. At 24hr post-treatment, mice were infected with HCMV TB40/E-infected fibroblasts (approximately 10^5^ PFU of cell-associated virus per mouse) via IP injection. A control group of engrafted mice was mock infected using uninfected fibroblasts. Virus was reactivated as previously described (35, 36).

### Quantitative PCR for viral genomes

DNA from CD34^+^ HPCs was extracted using the two-step TRIZOL (Thermofisher) method according to the manufacturer’s directions. Total DNA was analyzed in triplicate using TaqMan FastAdvanced PCR master mix (Applied Biosystems), and primer and probe for HCMV *UL141* and human β-globin as previously described (33, 35, 36). Copy number was quantified using a standard curve generated from purified HCMV BAC DNA and human β-globin-containing plasmid DNA, and data were normalized assuming two copies of β-globin per cell.

### Statistical analysis

Statistical analysis was performed using GraphPad Prism software (v10) for comparison between groups using student’s t-test, one-way or two-way analysis of variance (ANOVA) with Tukey’s post-hoc test or Bonferroni’s multiple comparison test as indicated. Values are expressed as mean +/- standard deviation or standard error of the mean, as indicated in the figure legends. Significance is highlighted with p<0.05.

## ACKNOWLEDGEMENTS

Research reported in this publication was supported by the National Institute of Health under Award Number T32AI170496.

## SUPPLEMENTAL FIGURE LEGENDS

**Figure S1.**
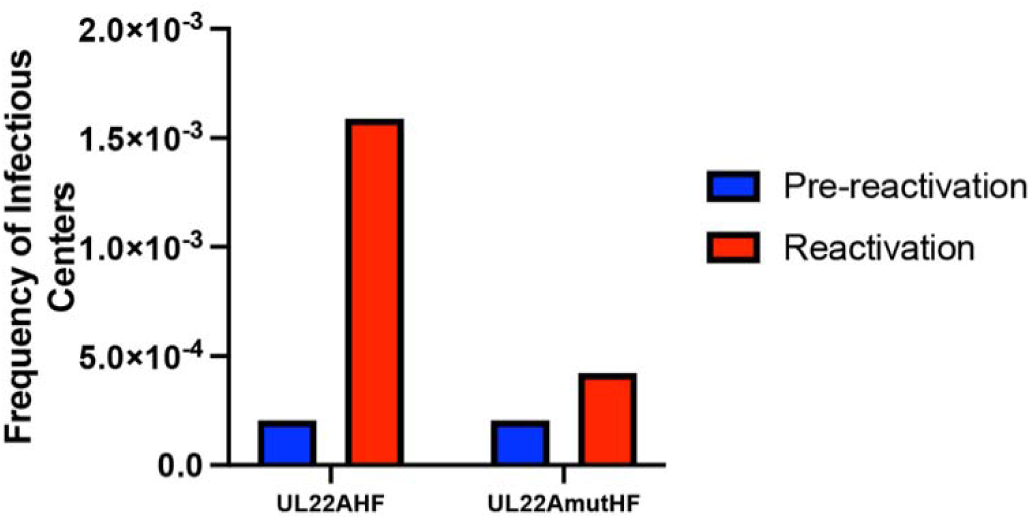
UL22A is necessary for reactivation from latency in primary fetal liver-derived CD34^+^ HPCs. Fetal liver-derived CD34^+^ HPCs were infected with TB40/E UL22AHF or UL22AmutHF at an MOI of 3 for 48 hours and then sorted using FACS for viable, CD34^+,^ GFP^+^ cells. Latency and reactivation experiments were performed as in Figure 2B.

**Supplemental Figure 2.**
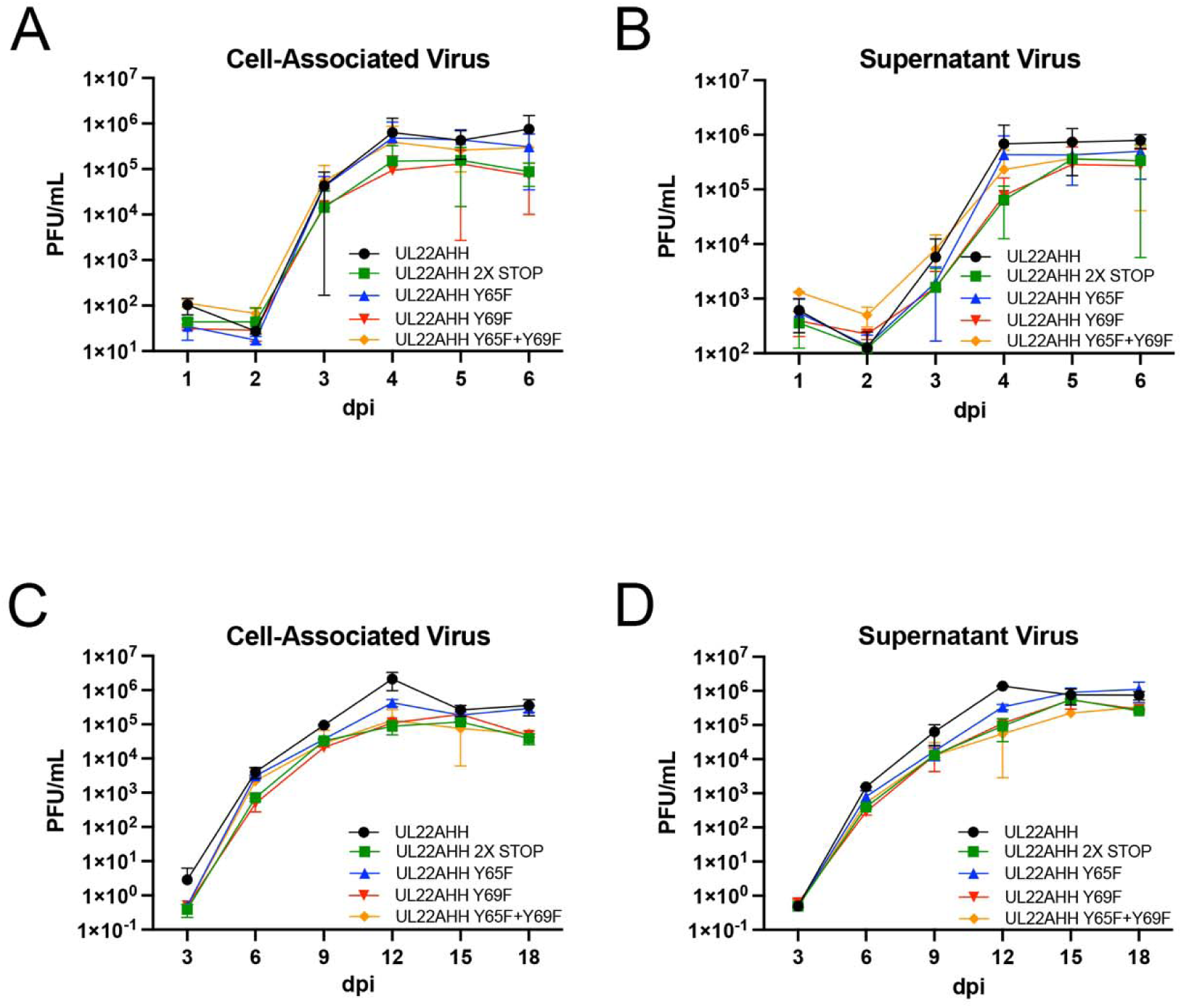
UL22A tyrosine residues Y65 and Y69 are not necessary for lytic replication. NHDFs were infected with TB40/E UL22AHH, UL22AHH 2X STOP, UL22AHH Y65F, UL22AHH Y69F, and UL22AHH Y65F+Y69F at an MOI of 3 for single step (A and B) or an MOI of 0.01 for multistep (C and D) growth curves. Cell-associated and supernatant virus were harvested at the indicated timepoints, and PFU/mL values were quantified. Data represents three independent experiments.

**Supplemental Figure 3.**
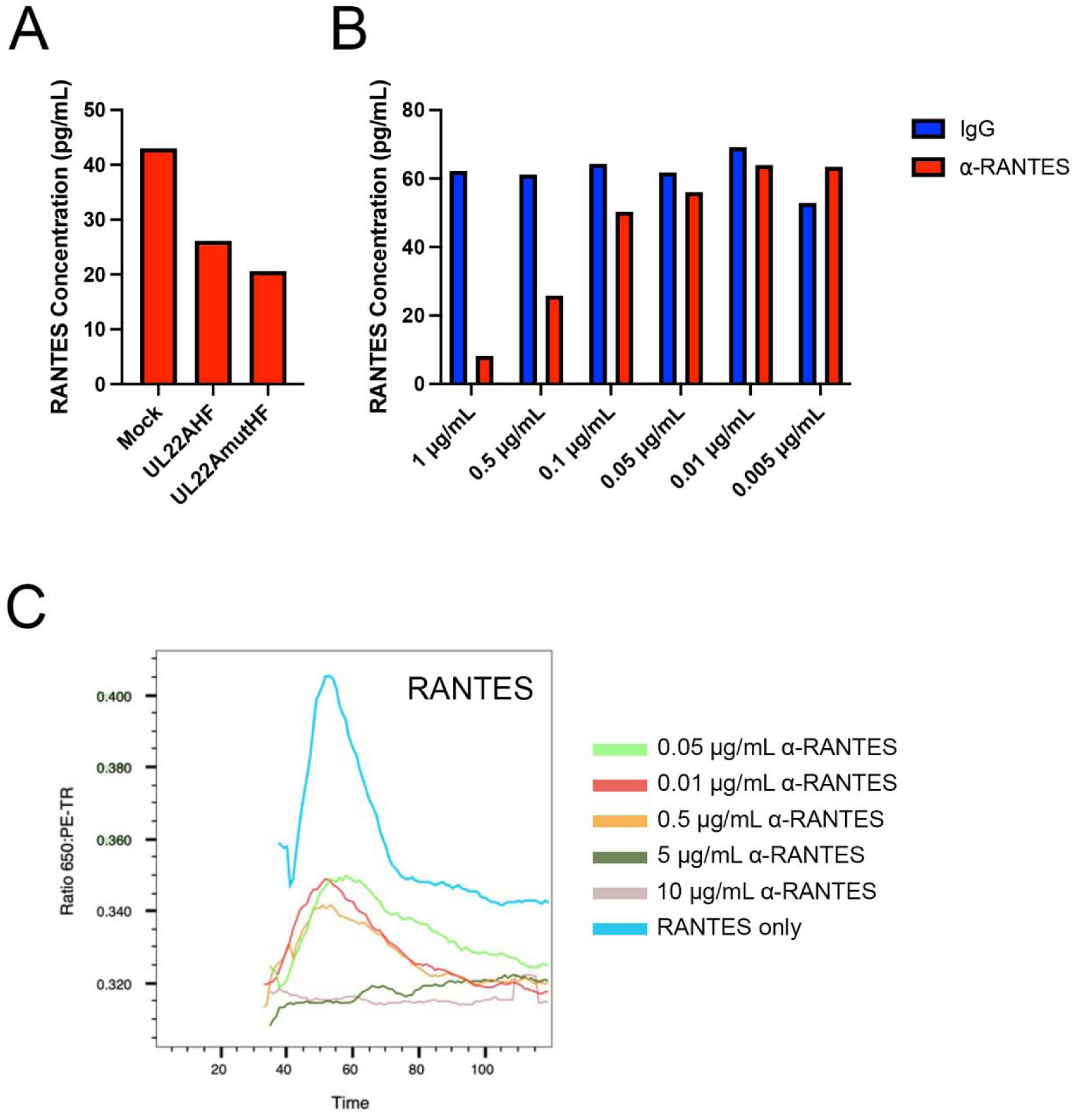
Validation of RANTES neutralization during latency using α-RANTES antibody. (A) hESC-derived CD34^+^ HPCs were Mock-infected, infected with TB40/E UL22AHF or UL22AmutHF at an MOI of 2 for 48 hours and then sorted using FACS for viable, CD34^+,^ GFP^+^ cells. Sorted cells were put into latency culture as in Figure 2B. At 14dpi, supernatants were taken and RANTES concentration was quantified using the Human CCL5/RANTES ELISA Kit (R&D Systems). (B) Control IgG or α-RANTES antibody at the indicated amounts were added to 50pg recombinant human RANTES for 2 hours, then RANTES was detected using the Human CCL5/RANTES ELISA Kit (R&D Systems). (C) CEM cells resuspended in 37°C HBSS with 1uM Fura Red, AM for viability and incubated for 30 minutes in a 37°C water bath. Recombinant RANTES and anti-RANTES were incubated at indicated concentrations for 30 minutes prior to assay. Background calcium flux was recorded for 20 seconds, sample was removed while recording, and RANTES (+/- anti-RANTES) was added to the cell suspension and placed back on the cytometer for recording. Events were recorded for 120 seconds.

**Supplemental Figure 4.**
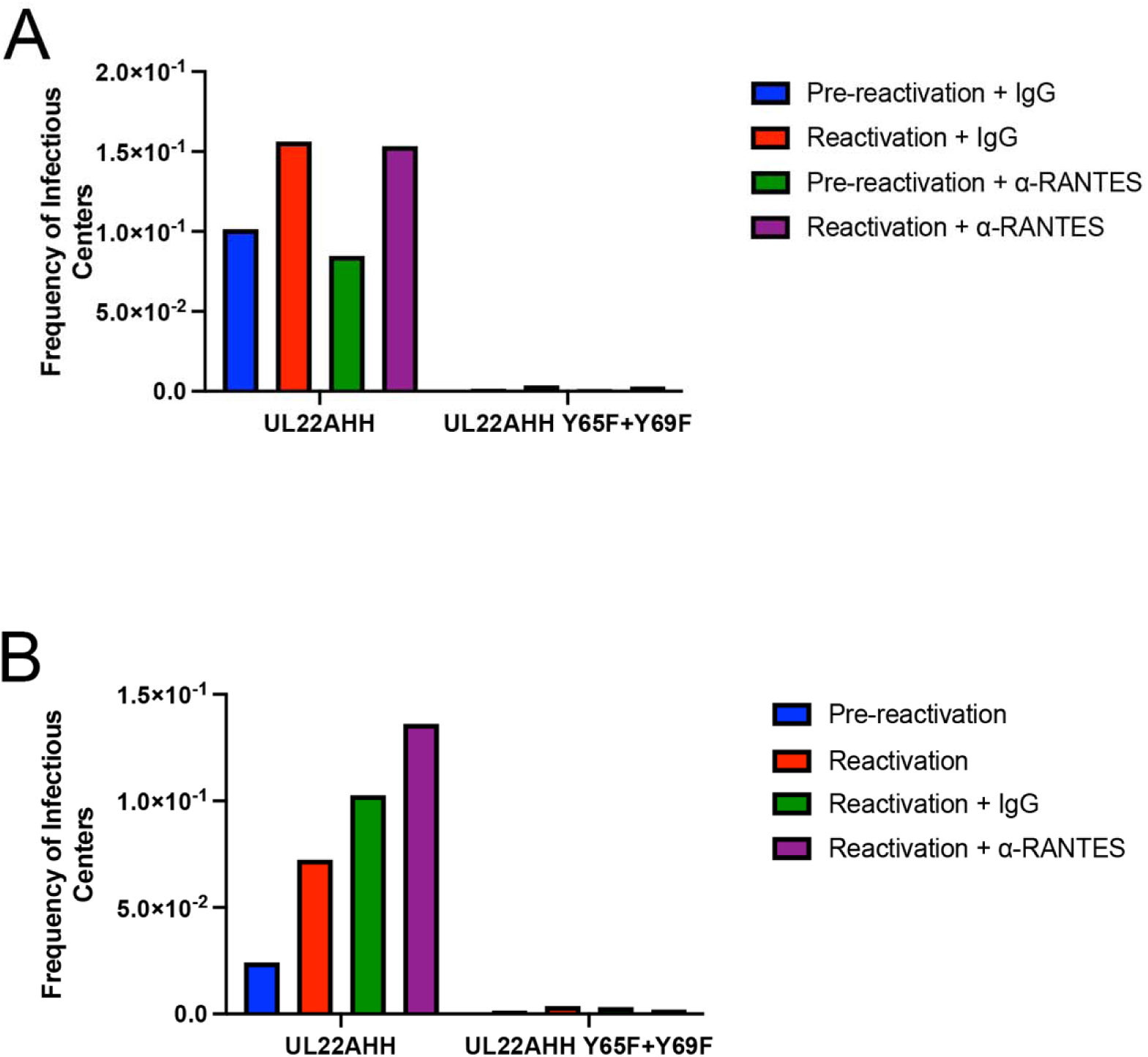
Adding 5ug/mL of anti-RANTES antibody during latency does not complement the UL22A mutant viruses. hESC-derived CD34^+^ HPCs were infected with TB40/E UL22AHH or UL22AHH Y65F+Y69F at an MOI of 2 for 48 hours and then sorted using FACS for viable, CD34^+,^ GFP^+^ cells. (A) Cells were treated with 5ug/mL of α-RANTES antibody or a control IgG and replenished again after 5 days while maintained in LTBMC in transwells above a murine stromal support to establish latent infection. (B) For 12 days, cells were maintained in LTBMC in transwells above a murine stromal support to establish latent infection. At 14 dpi, an equal number of cells were mechanically disrupted and seeded over a fibroblast monolayer to measure virus in latency culture (pre-reactivation) or cultured in cytokine-rich media containing 5ug/mL of α-RANTES antibody or a control IgG. Control IgG or RANTES neutralizing antibody were replenished after 1 week of reactivation. At 21 days post-plating, GFP^+^ wells were counted, and the frequency of infectious centers were determined by ELDA software.

